# An optimized tissue clearing protocol for rat brain labeling, imaging, and high throughput analysis

**DOI:** 10.1101/639674

**Authors:** Audrey Branch, Daniel Tward, Anna C Kolstad, Vyash Pulyadi, Joshua T Vogelstein, Zhuhao Wu, Michela Gallagher

**Affiliations:** Department of Psychological and Brain Sciences, Johns Hopkins University, Baltimore, USA; Departments of Computational Medicine and Neurology, Ahmanson-Lovelace Brain Mapping Center, University of California Los Angeles, USA; Department of Biomedical Engineering, Johns Hopkins University Baltimore, USA; Department of Cell, Developmental & Regenerative Biology, Icahn School of Medicine at Mount Sinai, New York, USA; Department of Neuroscience, Icahn School of Medicine at Mount Sinai, New York, USA

## Abstract

The advent of whole brain clearing and imaging methods extends the breadth and depth at which brain-wide neural populations and structures can be studied. However, these methods have yet to be applied to larger brains, such as the brains of the common laboratory rat, despite the importance of these models in behavioral neuroscience research. Here we introduce AdipoClear+, an optimized immunolabeling and clearing methodology for application to adult rat brain hemispheres, and validate its application through the testing of common antibodies and electrode tract visualization. In order to extend the accessibility of this methodology for general use, we have developed an open source platform for the registration of rat brain volumes to standard brain atlases for high throughput analysis.

## Introduction

Understanding the structure and function of the myriad of cells comprising the central nervous system requires precise methods for labeling and measuring intact neural cell populations and their connectional organization in circuits and networks. The development of brain clearing protocols in recent years has greatly enhanced our ability to visualize fluorescently tagged cell populations in the intact brain. Such methods have now been used to identify components of coactivated neural populations using immediate early genes (Renier et al., 2016), to delineate connectivity between brain regions (Ye et al., 2016), to assess three-dimensional transcriptional profiling (Wang et al., 2018), to demonstrate complex connections between the nervous system and peripheral tissues (Chi et al., 2018; Wang et al., 2018), and to illuminate circuit level connectivity (Lerner et al., 2015). However, these methods have yet to be applied broadly to larger animal models, as they are mostly optimized for mouse tissues and often for detecting transgenic reporters. For example, while clearing methods have been tested for laboratory rat brains (Stefaniuk et al., 2016), such applications often rely on the expression of fluorescent reporters. Rats have long been the model of choice for behavioral and cognitive neuroscience as they exhibit a broad and rich repertoire of quantifiable behaviors and their larger size assists in stereotaxic targeting of brain regions of interest. However, their larger size also presents a challenge in the application of brain clearing methodology, as these methods depend on the complete breakdown and/or removal of lipids, with subsequent refractory index matching throughout the tissue to achieve optimal transparency. Furthermore, there is a relative deficit of transgenic rat models, limiting the potential application of clearing methods such CLARITY and CUBIC (Chung et al., 2013; Susaki et al., 2015). Efforts to apply whole brain lightsheet imaging methods to rats models are further hampered by an absence of data processing methods applicable to the large volume datasets from cleared rat brain tissue imaging. To address these issues, we have optimized a brain clearing method for application to rat brain hemispheres, and developed a companion open source platform optimized for artifact removal and registration of the resulting lightsheet imaged brain data sets to a standard rat brain atlas for robust, automated, and reliable processing adaptable to regular laboratory settings.

Here, we successfully applied the iDISCO+ protocol (Renier et al., 2016) to rat brain hemispheres. This protocol was chosen for its compatibility with immunolabeling, minimal tissue shrinkage, its simplicity, and its low cost. Based on the recent advanced version, AdipoClear (Chi et al., 2018), we further optimized the protocol for rat brain lipid removal to ensure even clearing throughout an intact rat brain hemisphere and to facilitate later immunolabeling, while preserving brain 3D morphology for downstream automated data processing. We thus named the new protocol AdipoClear+, and tested it with immunolabeling against endogenous proteins and virally expressed reporters for applications. To maximize the application of AdipoClear+ method for use in rat models, we developed a registration method that can effectively process lightsheet imaging for artifact removal and volume image alignment to any MRI based rat brain atlas, which can be run using a standard desktop computer.

## Results

### AdipoClear+ delipidation and clearing for rat brain tissue

Rat brain tissue presents a challenge for whole brain clearing and imaging methodologies. The rat brain is larger than that of the mouse and contains a thicker myelination profile. For broad and convenient application to rat brain tissue, we reasoned that the protocol must be able to label endogenous or exogenous markers, as rat-based studies tend to rely on virally expressed or endogenous protein targets rather than endogenously expressed fluorescent markers. We began with the published iDISCO+ protocol (Renier et al., 2016) which is optimized for mouse brains with immunolabeling. Initial application of this protocol to adult rat brain tissue at 6 months of age resulted in incomplete clearing in the center of the tissue and slight discoloration, likely due to incomplete removal of lipids and oxidation, respectively (Fig. 1B). Given the larger size and/or thicker myelin, and older chronological age of our rat subjects relative to the mice for which the protocol was developed (Fig. 1D for size comparison) we speculated that rat brain tissue may require enhanced delipidation to fully clear the tissue, and thus sought to enhance the effectiveness of the lipid removal steps.

**Figure 1:**
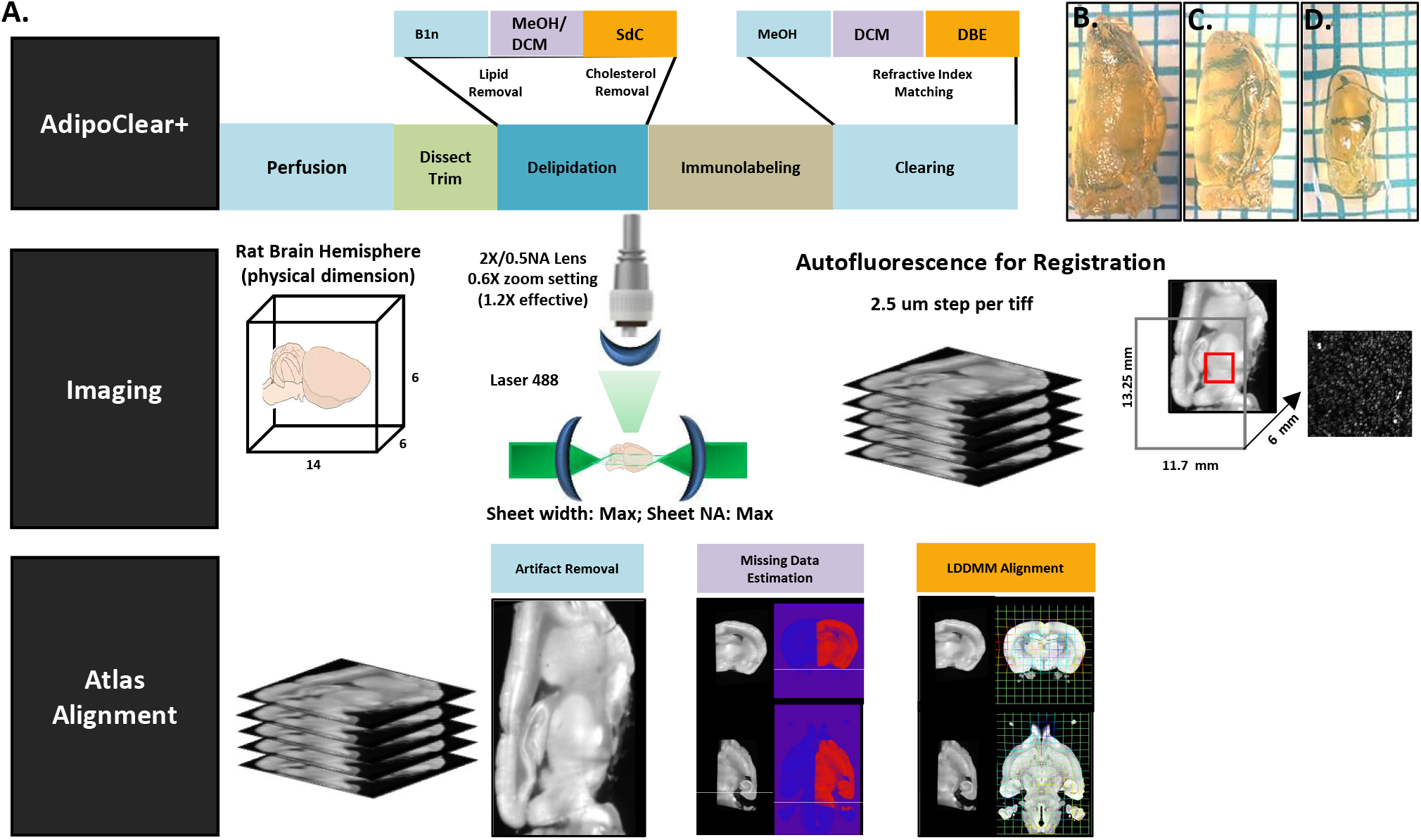
**A:** Overview of the rat brain optimized clearing, imaging, and registration pipeline. **B:** Rat brain hemisphere cleared with original iDISCO+ protocol retains opaque center. **C:** Rat brain and **D:** mouse brain hemisphere cleared with AdipoClear+ protocol.

Toward this goal, we adapted the delipidation methods of the AdipoClear protocol (Chi et al., 2018), which was developed to clear and immunolabel lipid dense adipose tissue without damaging tissue structure. To improve clearing of the deep myelin region, we designed an additional washing buffer with methyl-β-cyclodextrin (MβCD). MβCD offers high affinity extraction of cholesterol, the major component of dense myelin lipids, to ensure complete delipidation for optimal clearing and immunolabeling. A schematic of this modified protocol, AdipoClear+, with relevant additions and substitutions is shown in Fig. 1A, alongside a rat brain hemisphere cleared with this new protocol (Fig. 1C), as compared to results of the previous version of the protocol (Fig. 1B). A full outline of protocol steps and recommendations can be found in Supplemental Figure 1.

In order to ensure that the AdipoClear+ protocol is fully compatible with rat brain immunolabeling, we tested several different antibodies which have previously been shown to be compatible with iDISCO+. Brains were imaged using a LaVision BioTec Ultramicroscope with a 2X variable zoom lens on 0.63X magnification, which allows for an entire rat brain hemisphere to be imaged in either sagittal or horizontal orientation. To achieve even illumination across the tissue, both lasers were used for all scans and the maximum lightsheet width and NA were used. Figure 2 shows representative results of staining in a rat brain with several common antibodies. Images are shown for anti-tyrosine hydroxylase staining throughout the z-extent of the tissue, demonstrating no loss of image resolution with increasing z-depth (Fig. 2A-E, A1-E1). Previously validated antibodies including parvalbumin (Fig. 2F), and GFP and RFP (Fig. 2G) are also compatible with the protocol.

**Figure 2:**
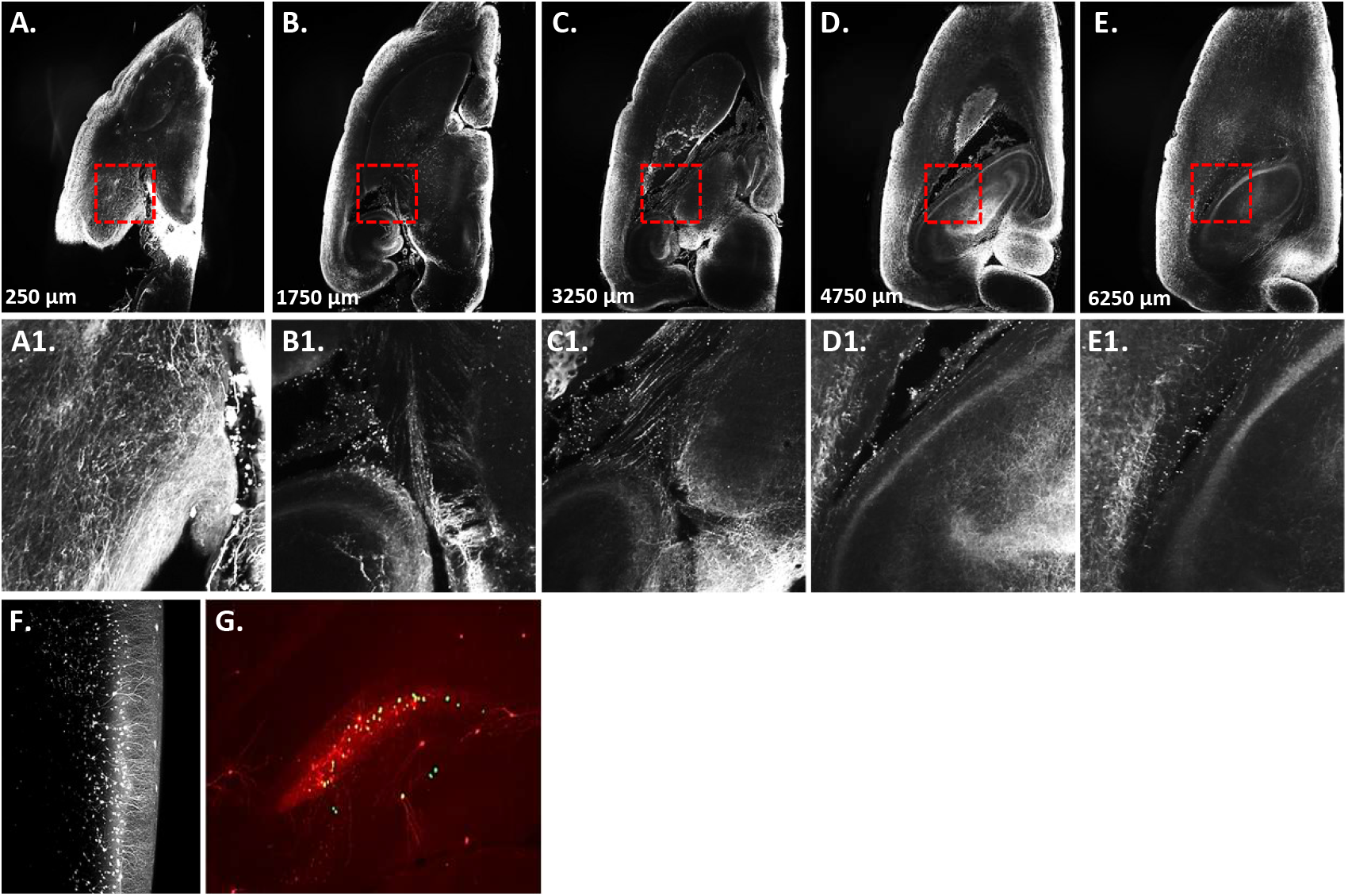
Validation of immunolabeling for rat brain optimized AdipoClear+. **A-E:** Anti-tyrosine hydroxylase staining, whole brain. Images are z-projections (50 tiffs); approximate depth in tissue indicated. **A1-E1:** Zoom of box outlined in A-E demonstrating showing resolution of signal at increasing tissue depth. **F:** Immunolabeling of parvalbumin in cortex. **G:** Immunolabeling of virally expressed GFP and RFP in hippocampus.

### Application to whole brain activity mapping and electrode tract verification

The laboratory rat is a staple model for behavioral neurobiology. However, it remains challenging to link neural activity to behavioral performance. Common methods of identifying coactive populations of cells include immediate early gene (IEG) mapping and electrophysiological recording. The use of IEGs, such as cFos, whose expression levels are reflective of recent changes in neuronal activity allows for identification of coactivated cell populations on a brain wide level. While the temporal resolution of IEGs is low with their expression lagging from several minutes to hours after high activity firing (Barnes, 2015), the spatial resolution of this approach enables unbiased identification of brain regions and cell populations that underly behavioral activation. However, slice-based approaches to quantification of IEG expression are arduous, as brains are sliced and imaged individually then manually annotated, typically only in predetermined brain regions. Whole brain clearing and imaging provides a potential avenue for capturing global activity profiles with IEG labeling (Renier et al., 2016), but such methods have yet to be applied to the laboratory rat. To address this, we tested our protocol for compatibility with cFos labeling (Fig. 3A-E, A1-E1). The improved clearing performance of the optimized protocol makes it possible to image cFos positive cells throughout the extent of the intact hemisphere, allowing for brain wide mapping of coactive cells via automatic cell detection and quantification methods, such as ClearMap (Renier et al., 2016).

**Figure 3:**
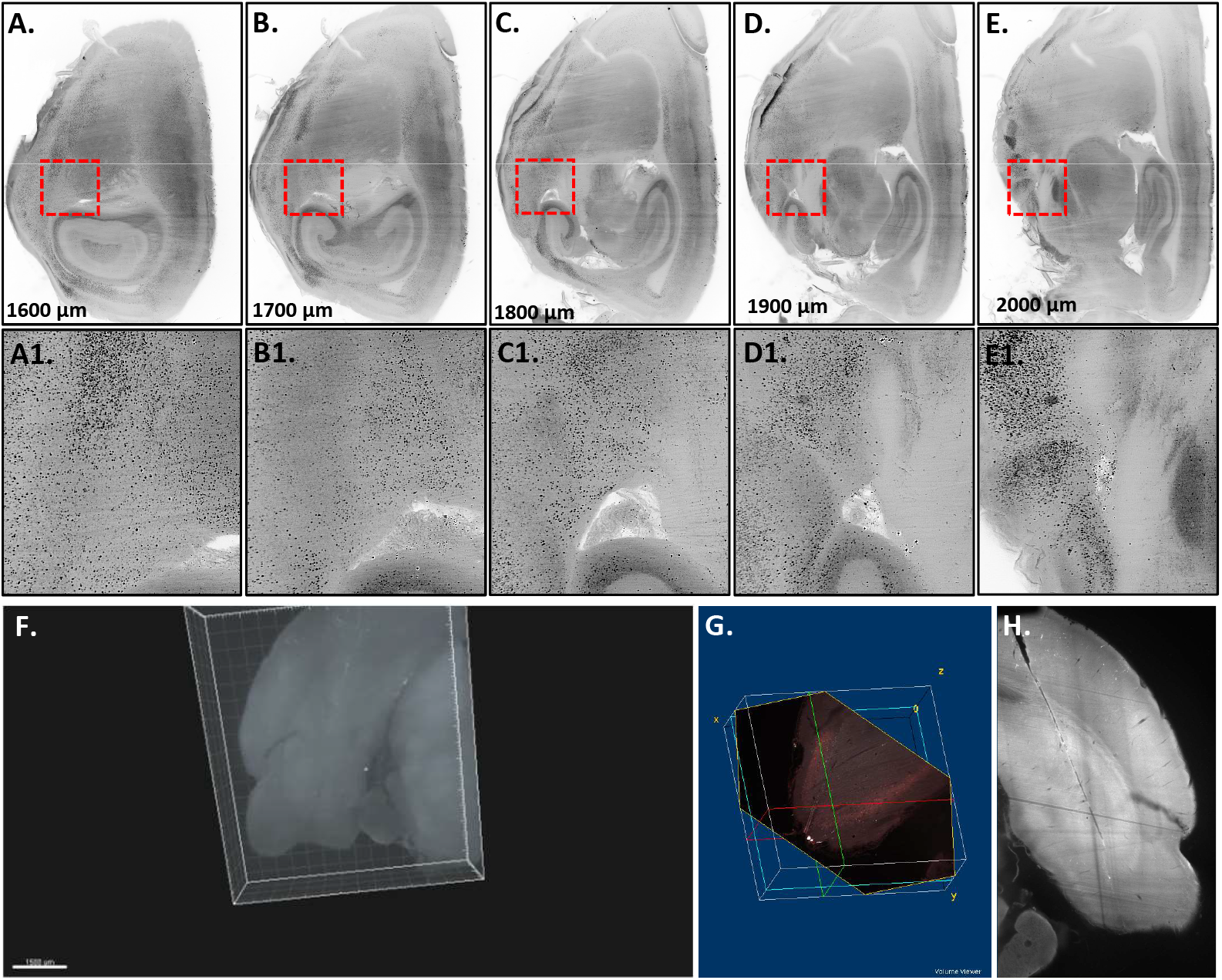
Global mapping of activity and electrode tract imaging. **A-E:** cFos immunolabeling of rat brain hemisphere. Images are z-projections (100 tiffs); approximate depth in tissue indicated. **A1-E1:** Zoom of boxed region, displaying resolution of cFos positive cells from corresponding images. **F:** 3D volume rendering of AdipoClear+ cleared rat brain showing electrode track of two shank silicon probe. **G:** ImageJ 3D Volume Viewer can be used to obliquely reslice a stack containing an electrode track, to generate **H:** Coronally imaged stack resliced with a rotation of 18 degrees in x, 5 degrees in y to clearly show the of one electrode shank.

Electrophysiological recording makes up for the temporal imprecision of IEGs, providing instantaneous readouts of the electrical activity of cells. The recent development of high density silicon probes (Jun 2017) has dramatically increased the number of cells that can be recorded simultaneously from the level of dozens to hundreds and promises to extend our understanding of how neurons act in concert at a population and network level (Steinmetz, Koch, Harris, & Carandini, 2018). However, in order to be meaningfully interpreted, it is essential that electrode placement can be precisely verified to determine the location of recorded cells. Electrode tract verification is currently performed by slicing, serial imaging, and painstaking manual reconstruction of individual tissue sections. Here, we tested the ability of AdipoClear+ clearing and lightsheet imaging to allow these tracts to be identified in a 3D brain volume. For this, animals were implanted with dual shank 64 channel silicon probes (Cambridge Neurotech) targeting the right lateral entorhinal cortex (LEC). Over the course of several days, behavioral recording experiments were performed and the probes were lowered 400 μm per day until they exited the brain. Animals were then perfused, probes were fully retracted, and brains were extracted and processed. Given that the electrode tract was expected to span only the anterior portion of the right brain hemisphere, brains were trimmed to an appropriate size that contained the electrode tract and anatomically relevant surrounding tissue. Given the smaller size of the tissue, AdipoClear+ delipidation and permeabilization steps were performed with shortened incubation steps. We took advantage of the inherent autofluorescence of AdipoClear+ cleared tissue in the 488 nm wavelength and generated a lightsheet image stack containing the electrode tract (Fig. 3F). While the tract is visible as a white line within a 3D rendered volume (Imaris), more precise identification of the path of travel and termination point can be obtained by reslicing the image stack via freely available Volume Viewer plugin of ImageJ (Fig. 3G, H), which allows for a tiff stack to be resliced obliquely along user selected x, y, and z planes. Together, these approaches make it possible to precisely locate the tract of an electrode in an intact rat brain, minimizing the chance of error in locating tracts due to slicing and mounting artifacts and reducing the manual effort of tissue processing and tract reconstruction.

### Registration of cleared brain image data to rat brain atlas

Extracting meaningful information from cleared intact brain tissue typically involves quantifying the distribution of labeled cells within anatomically meaningful regions. Our imaging of rat brain hemispheres generated data sets with ~2500 tiffs per channel in sagittal orientation and ~3500 in horizontal and are therefore too large to be manually annotated. While some methods do exist for automatically registering lightsheet imaged brains to atlases, they are typically designed for use with mouse tissue (Renier et al., 2016) or for application to aqueous based clearing methods (K. S. Kutten et al., 2017; K. S. Kutten, Vogelstein, J.T., Charon, N., Ye, L., Deisseroth, K., Miller, M.I., 2016; Susaki et al., 2015), which have a very different intensity profile than tissue cleared with iDISCO family protocols. We therefore sought to develop an automated annotation method optimized for AdipoClear+ cleared rat brain tissue. The most common approach to this form of annotation involves estimating a nonlinear registration between observed data and a well characterized atlas, then using the transformation to transfer annotations from the atlas onto the observed dataset. A variety of methods exist for registering medical image data to standard brain atlases (Klein et al., 2009). Given the similarity of the data structure between MRI and cleared brain imaging, efforts have been made to apply these methods for the registration of cleared brain image sets to atlases, using a combination of rigid and/or affine deformations of the sample data to align it with the atlas space (Renier et al., 2016; Susaki et al., 2015).

However, light sheet microscopy images of cleared tissue present challenges not seen in human MRI. Specifically, the major challenges needing to be addressed are image registration in the presence of missing or damaged tissue and variable image contrast. We use an image registration framework developed originally for digital pathology (Tward, 2019) that functions robustly in the presence of these issues. In this framework, contrast differences between atlas images and target images are estimated jointly with image registration parameters. The location of missing and damaged tissue or other artifacts, is also estimated jointly in an iterative fashion using the Expectation Maximization algorithm (Dempster, 1977).

Our starting data set was obtained by imaging a rat brain hemisphere positioned in sagittal orientation using a 2X variable zoom objective at a 0.63X zoom setting (1.2X effective magnification). Using the 488 nm laser and imaging throughout the z extent of the hemisphere with a 2.5 μm step size resulted in a data set of 2500 individual tiffs representing the autofluorescent intensity profile of major anatomical structures spanning the entirety of the hemisphere. We choose the Waxholm MRI atlas (Kjonigsen, Lillehaug, Bjaalie, Witter, & Leergaard, 2015; Papp, Leergaard, Calabrese, Johnson, & Bjaalie, 2014) for initial testing of the alignment of this data set with this methodology. An example result illustrating image correction steps is shown in Fig.4, and image annotation through registration is shown in Fig. 5. Panel C shows anatomical structures defined in the atlas as colors, overlayed on the grayscale atlas image. A good alignment can readily be observed for subcortical gray matter structures, as well as for cortical boundaries. Panel D shows a comparison between atlas and target image intensities, where yellow indicates relatively good agreement between the images.

**Figure 5.**
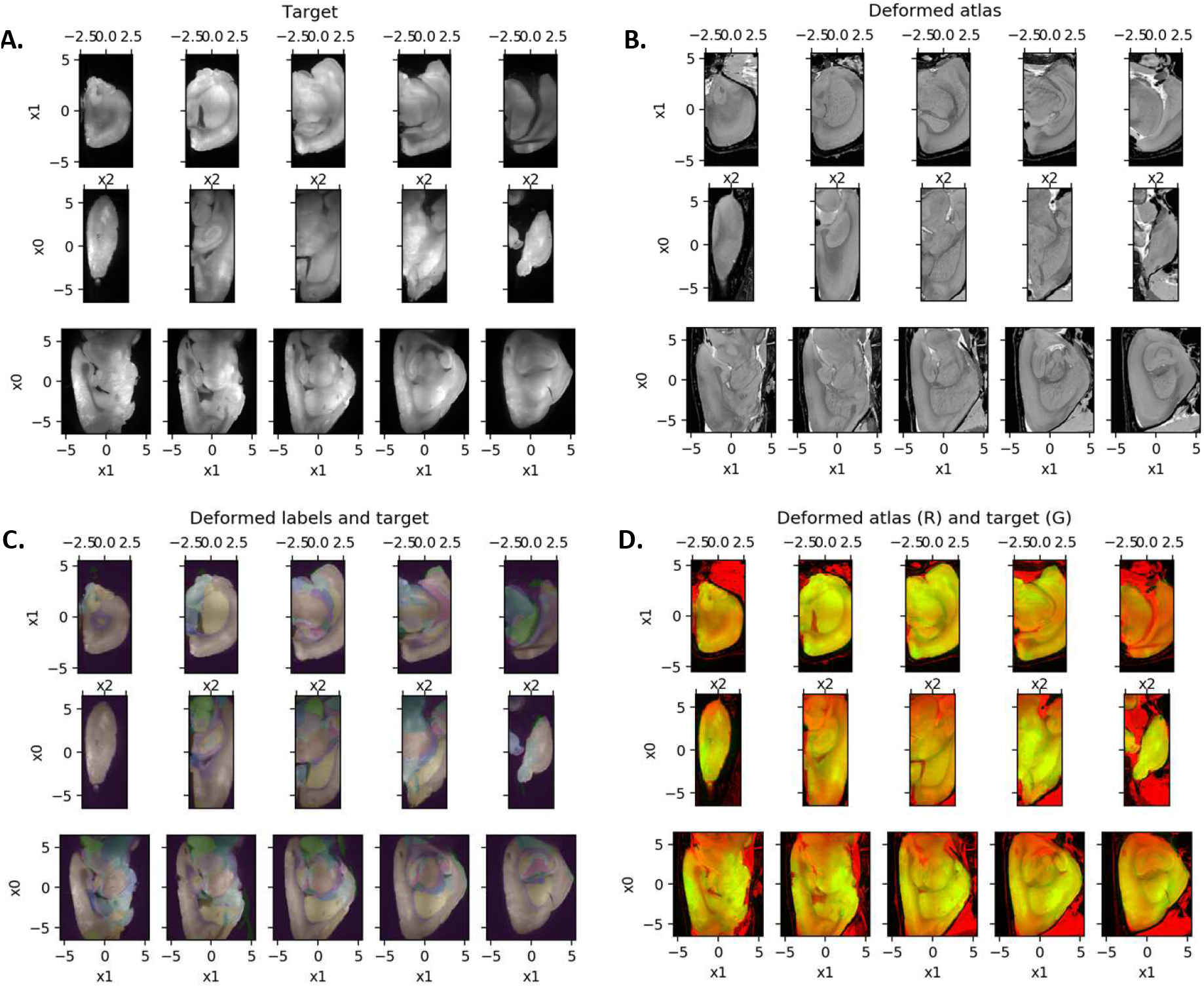
**A:** Target image is aligned with a deformed atlas image (**B**). Annotation labels are showed overlaid on target image (**C**), and both images are shown as red and green channels (**D**) yielding yellow where image intensities are similar. All images are shown with 5 slices in 3 planes.

Image registration typically involves many parameter choices that result in a barrier to use for neuroscientists. To improve accessibility of our approach, we have made several examples publicly available through github [https://github.com/dtward/image_lddmm_tensorflow], using Jupyter notebooks [https://jupyter.org/] as an intuitive web browser based interface. See Example_iDISCO_rat_waxholm.ipynb in the repository for one example from which the figures in this paper are generated. All necessary parameter choices have been made and justified within the Jupyter notebook. This approach can be extended to a variety of other model organisms and humans to support similar subject of future work for broad applications.

### Discussion

Unbiased assessment of neuronal populations in intact brains is a valuable exploratory tool, but remains limited in its application. We have developed a brain clearing methodology that is capable of evenly immunolabeling and clearing an adult rat brain hemisphere using adult Long Evans rats. We began by applying the solvent based iDISCO+ (Renier et al., 2016) methodology, which offers several advantages for application to the larger brain of the common Long Evans outbred rat strain. AdipoClear+ delipidation and clearing causes minimal distortion of tissue size as compared to aqueous based protocols which tend to enlarge samples (Wan et al., 2018) and was developed to optimize the penetration of antibodies for immunolabeling of endogenous proteins. In addition, the inherent autofluorescence present in the 488 nm wavelength spectrum of AdipoClear+ cleared brains provides highly detailed structural information which can be used to register resulting image data sets to standard MRI based brain atlases. Further, we have extended the application of this methodology to allow for whole brain activity mapping with cFos, and for 3D validation of electrode placement. Taken together, this rat brain clearing and imaging platform makes efficient whole mount analysis for broad brain research using the common laboratory rat.

## Methods

### Animals

Adult male Long-Evans rats were obtained at 3-4 months of age from Charles River Laboratories (Raleigh, NC) and housed in a vivarium at Johns Hopkins University. All rats were individually housed at 25°C and maintained on a 12 hr light/dark cycle. Food and water were provided ad libitum until indicated otherwise. The rats were examined for health and pathogen-free status throughout the study, as well as necropsies at the time of sacrifice. All procedures were approved by the Johns Hopkins University Institutional Animal Care and Use Committee in accordance with the National Institutes of Health directive.

### Electrode surgery

For electrode tract experiments, 2 male Long Evans rats underwent implantation of multi-channel silicon probes under sterile conditions similar to chronic implantation procedure described previously (Deshmukh 2011). Under surgical anesthesia, animals were implanted with a 2 shank, 64 channel silicon probe (Cambridge Neurotech, cat#: H6-ASSY-156) directed at the lateral entorhinal cortex (LEC) with a 15 degree angle relative to the bregma/lambda horizonal reference plane with tips pointing in the lateral direction. The probes were inserted 4.6 to 5.0 mm lateral to the midline and 7.5-7.8 mm posterior to bregma with an initial depth of 5.5 mm. The implant was secured to the skull with self-adhesive resin (3M, cat#: 56830) and dental cement. Over the course of behavioral experiments, probes were lowered 400 μm per day until they exited the brain. Following recording, animals were heavily anesthetized with an overdose of isofluorane and intracardiac perfusion and fixation was performed with PBS+NaN3 followed by 4% PFA. Brains were fixed overnight in 4% PFA, then probes were fully retracted prior to brain extraction.

### Sample collection

Mature adult male rats (6-10 months old) were heavily anesthetized with an overdose of isofluorane and intracardiac perfusion and fixation was performed with PBS with 0.02% NaN3 followed by 4% PFA. All samples were post-fixed in 4% PFA at 4°C overnight. Fixed samples were then washed in PBS+NaN3 for 1 hour three times. Prior to beginning the protocol, brains were cut down the midline. As rat brains have thicker meningeal tissue relative to mice, this tissue was carefully removed with tweezers to allow for better diffusion of solutions and antibodies into the brain.

### AdipoClear+ Rat Protocol

| Reagent | Reference |
| --- | --- |
| PBS 10X | Ambion AM9624 |
| ParaFormAldehyde 16% (PFA) | EMS 15710-S |
| Triton X-100 | Sigma X100-500ML |
| Tween-20 | Sigma P9416-100ML |
| Glycine | Sigma G7126-500G |
| Sodium Azide (NaN<sub>3</sub>) | 58032-100G |
| 10N Sodium Hydroxide (NaOH) | Fisher SS255-1 |
| Heparin | Sigma H3393-50KU |
| DMSO | Fisher D128-4 |
| Methanol (MeOH) | Fisher A412SK-4 |
| Hydrogen Peroxide 30% (H<sub>2</sub>O<sub>2</sub>) | Sigma 216763-100ML |
| DiChloroMethane (DCM) | Sigma 270997-12X100ML |
| DiBenzylEther (DBE) | Sigma 108014-1KG or 3KG |
| methyl-β-cyclodextrin (MβCD) | Sigma 332615 |

### Delipidation and Permeabilization

For immunolabeling, fixed rat brain hemispheres were washed in B1n buffer (500 mL H2O, 0.5% TritonX, 2% glycine, 50 ul 10M NaOH, 0.02% NaN3) 2 hr, 4 hr, and overnight. Samples were then washed/dehydrated in a 20%, 40%, 60%, 80% MeOH/B1n gradient for 1 hr each. Samples were then treated with 100% dichloromethane (DCM) for 30 minutes three times, then overnight. DCM was washed out with 100% MeOH for 1 hr, 3 hr, and overnight and rehydrated with a reverse gradient of MeOH/B1n 80%, 60%, 40%, 20% for 1 hr each, and 100% B1n for 1 hr then overnight. This was followed by washes with SdC buffer for 4 hr followed by 2 × 24 hr, and 2 × 48 hr. Finally, samples were treated with a 5% DMSO/0.3M Glycine/PTxwH solution at 37°C for 6h and overnight, then washed in PTwH for 1.5 hr, 3 hr, and overnight (PTxwH: 500 mL PBS, 0.5% TritonX, 0.25% Tween-20, 50 uL heparin stock solution (20 mg/mL), 0.02% NaN3). Except otherwise noted, all steps were performed with shaking at 150 RPM at RT. For electrode tract identification, brain hemispheres were trimmed to a smaller size and a minimal protocol was devised for delipidation and permeabilization. The steps were performed as above, but SdC buffer incubation was to 1.5 hr, 3 hr, and overnight, and immunolabeling steps were not performed. For step by step protocol instructions, see supplemental materials.

### Immunolabeling

Samples were incubated in primary antibody solution (primary antibody in 5% DMSO/0.3M Glycine/PTxwH) for 8 days. After primary antibody incubation, samples were washed in PTwH for 3 hr, 4 hr, overnight, and 4 × 24 hr to wash out residual primary antibody. Samples were then incubated for 8 days in secondary antibody solution (secondary antibody, PTwH), then washed PTwH for 3 hr, 4 hr, overnight, and 4 × 24 hr to wash out residual secondary antibody. In this study, we tested the following primary antibodies: rabbit anti-cFos (Cell Signaling, cat# 2250; 1:200), rabbit anti-parvalbumin (Swant, cat#: PV27, 1:200), sheep anti-tyrosine hydroxylase (Millipore, AB1542, 1:200), chicken anti-GFP (Aves, cat#: GFP-1020, 1:1000), and rabbit anti-RFP (Rockland, cat.#: 600-401-379, 1:1000). Secondary antibodies were used at the same concentration as their respective primary antibodies and included donkey anti-rabbit 568 (ThermoFisher, cat# A10042), donkey anti-chicken 647 (Millipore, cat# AP194SA6) or donkey anti-sheep 647 (ThermoFisher, cat# A-21448).

### Tissue Clearing

Samples were post-fixed in 2% PFA overnight at 4°C, then dehydrated in 20%, 40%, 60%, 80%, 100% H2O/MeOH series for 1 hr each at RT, followed by 100% MeOH overnight. Following dehydration, samples were washed in 100% MeOH 3 hr, 1:2 MeOH/DCM 2 hr, 100% DCM 1 hr, 3 hr, and overnight, followed by clearing with DBE. Samples were stored in the dark until imaging.

### Lightsheet imaging

All samples were imaged on a lightsheet microscope (Ultramicroscope II, LaVision Biotec) equipped with a 2X (low magnification, whole brain) or 4X (high magnification) lens and a sCMOs camera (Andor Neo). Images were acquired with the ImspectorPro software (LaVision BioTec). For imaging, samples were clamped in place in the sample holder in sagittal orientation (midline up), or horizontal orientation (dorsal up) and placed in an imaging reservoir filled with DBE and illuminated from the side by the laser light sheet. For low magnification scanning of whole hemispheres for atlas registration, samples were scanned with the 488 (filter: 575/40), 561 (filter: 620/60), or 640nm (filter: 680/30) laser channels with a step size of 2.5 μm for the 2X objective on 0.63X zoom setting (1.2X effective magnification) with a sheet aperture of 0.1. For imaging of cFos labeled brains, the 568 channel was imaged using the continuous light sheet scanning method with the included contrast blending algorithm to improve resolution and obtain even imaging (Renier et al., 2016).

### Image processing

Several preprocessing steps were used before registration. First, streak correction was performed by zeroing out signal at −10.5 degrees, 0 degrees, and 10 degrees in the Fourier domain as shown in Fig. 4. These artifacts are the result of shadowing when tissue boundaries are parallel to the light source. Second, slice to slice intensity variations were corrected such that the mean signal changed smoothly across slices. To accomplish this, mean intensity was computed as a function of slice number, and the resulting signal was smoothed using a Gaussian kernel with standard deviation 33 slices. Each slice intensity was then normalized by dividing by its mean intensity, and multiplied by its smoothed intensity. Third, 3D images were inhomogeneity corrected using the N4 algorithm implemented in simple ITK (Tustison et al., 2010). Fourth, images were downsampled by averaging neighboring voxels, to a resolution of 0.077 mm. This corresponds approximately to the resolution of the Waxholm MRI atlas after a factor of 2 downsampling.

**Figure 4:**
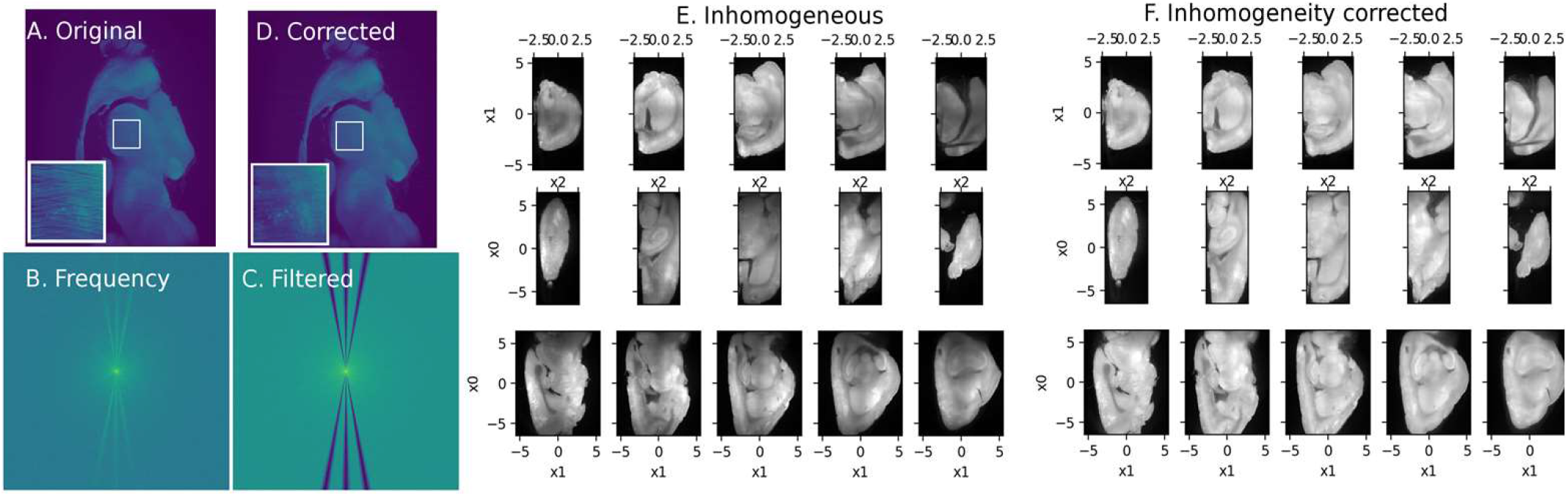
Imaging artifact correction. **A-D**: Streak correction. Left: before filtering, right: after filtering. Top: spatial domain, bottom: frequency domain. **E, F:** Nonuniformity correction. Nonuniform image (**E**) is corrected (**F**) using a spatially smooth multiplicative transform. Axes x0, x1, and x2 have units in mm.

Images for figures were generated using ImageJ (NIH, http://imagej.nih.gov/ij/). For signal channels, background subtraction was performed using a 50 μm rolling ball radius. For figures showing autofluorescent channels, no image modification was performed other than those discussed in each figure. Imaris (Bitplane, http://www.bitplane.com/imaris/imaris) was used for the video of 3D volume rendering of electrode implanted brain in Figure 3.

### Registration

We perform deformable image registration between the autofluorescence channel *J*: *X* ⊂ R^3^ → R (this notation means “J is a function that maps points in some subset of 3D space to real numbers”) and the Waxholm T2* rat atlas *I*: *X* → R. Autofluorescence has been successfully used for registration in similar modalities such as CLARITY (K. S. Kutten, Vogelstein, J.T., Charon, N., Ye, L., Deisseroth, K., Miller, M.I., 2016).

We construct spatial transformations *ϕ*: *X* → *X* using the Large Deformation Diffeomorphic Metric Mapping model (Beg M.F, 2005) solving a variational problem to minimize the weighted sum of a regularization cost and a matching cost. The diffeomorphic property (smooth, 1 to 1, with smooth inverse) is ensured by defining *ϕ* as the flow of a smooth velocity field *v_t_*: *X* → R^3^ for *t* ∈ [0,1]:

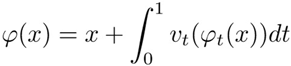

Sufficient smoothness is ensured by defining a regularization function via

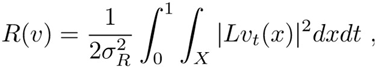

where *L* is the differential operator *L* = (*id* − *a*^2^Δ)^*p*^ (Dupuis P., 1998) for *a* a smoothness scale (with units of length), Δ the Laplacian 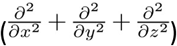, and *p* ≥ 2 power. We choose *a* =1 mm voxels (double check!) and *p* = 2. The parameter *σ_R_*^2^ is a weighting factor.

To choose an appropriate matching cost, we must overcome several important challenges intrinsic to this data. The image intensity profile differs from the MRI atlas, and tissue is missing due to cutting and limited field of view and artifacts are present.

We overcome these challenges simultaneously using estimation within a generative statistical model, rather than proposing a series of steps in a pipeline. Instead of using a similarity function such as normalized cross correlation or mutual information, we predict *J* in terms of 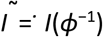 and use a simple Gaussian white noise model as a cost (sum of square error).

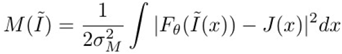

Where 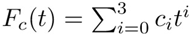 is a cubic polynomial which predicts the intensities in *J* from those in *I*^~^. The factor *σ_M_*^2^ is a weighting factor, chosen to be approximately equal to the variance of the noise in our Gaussian white noise model.

Using a white noise model allows us to accommodate missing data using the expectation maximization algorithm, as discussed in (Tward, 2019). In each *E* step we compute posterior probabilities that tissue falls into one of two classes: some location in the atlas image ((M)atching), or (A)rtifact and missing tissue. The data likelihood under the first class uses the model described above, while the latter is thought of as constant intensity (unknown mean *c_A_*) with Gaussian white noise of variance *σ_A_*^2^. We choose *σ_A_*^2^ = 10*σ_M_*^2^. Let 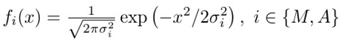 be a zero mean Gaussian CDF, the posteriors for the first class is

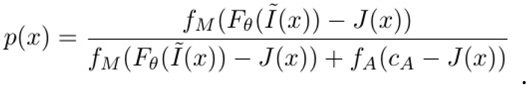

In the *M* step each unknown parameter is estimated by minimizing our cost function weighted by the posterior above. The mean *c_A_* can be found exactly via a weighted average. The *ϑ* can be found exactly by solving a linear equation (weighted least squares). In both cases the weights at each voxel are given by p(x). The *v_t_* is estimated iteratively using weighted LDDMM, as described in (Tward, 2019).

EM iterations result in a monotonically increasing complete data likelihood (Dempster, 1977). We note that this approach to missing data is only possible because our cost function represents a negative log likelihood.

This procedure was implemented in python tensorflow [https://www.tensorflow.org/], handling high performance computing requirements such as parallelizing across multiple cores or GPU.

Working examples are prepared through Jupyter notebooks (jupyter.org) and made publicly available through neurodata.io/reg.

### Registration Interface

An example Jupyter notebook illustrating our registration procedure can be found here: https://github.com/dtward/image_lddmm_tensorflow/blob/master/Example_iDISCO_rat_waxholm.ipynb. The registration procedure includes preprocessing, followed by registration at 2 different resolutions, using the lower resolution result as an initial guess for unknown parameters at higher resolution. Users can download this notebook and simply replace filenames with their data.

## Supporting information

Supplemental protocol

