## Supplemental protocol for "An optimized tissue clearing protocol for rat brain labeling, imaging, and high throughput analysis"

### Slide 1
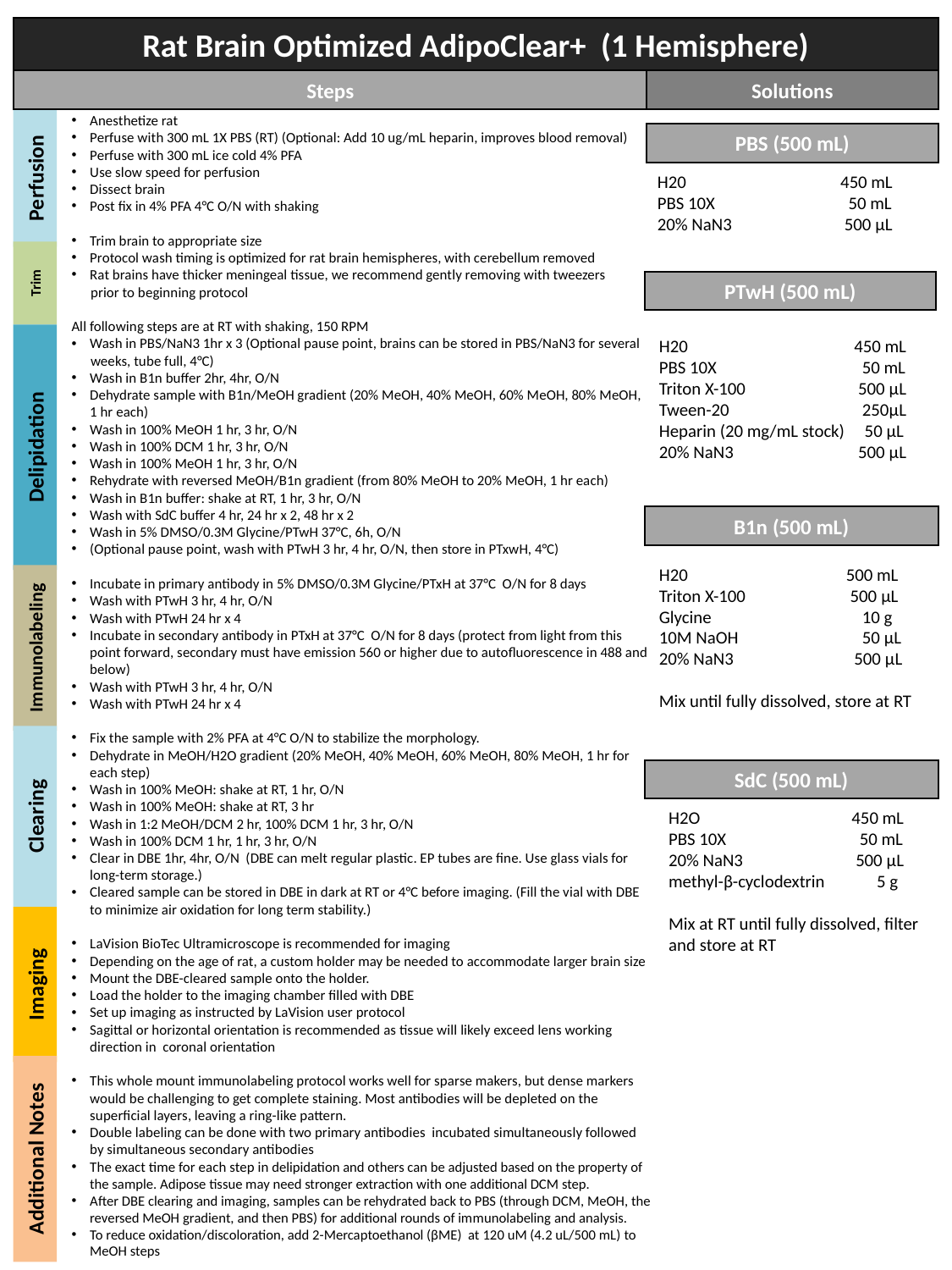

Rat Brain Optimized AdipoClear+ (1 Hemisphere)
Steps
Solutions
Anesthetize rat
Perfuse with 300 mL 1X PBS (RT) (Optional: Add 10 ug/mL heparin, improves blood removal)
Perfuse with 300 mL ice cold 4% PFA
Use slow speed for perfusion
Dissect brain
Post fix in 4% PFA 4°C O/N with shaking
Trim brain to appropriate size
Protocol wash timing is optimized for rat brain hemispheres, with cerebellum removed
Rat brains have thicker meningeal tissue, we recommend gently removing with tweezers
 prior to beginning protocol
All following steps are at RT with shaking, 150 RPM
Wash in PBS/NaN3 1hr x 3 (Optional pause point, brains can be stored in PBS/NaN3 for several
 weeks, tube full, 4°C)
Wash in B1n buffer 2hr, 4hr, O/N
Dehydrate sample with B1n/MeOH gradient (20% MeOH, 40% MeOH, 60% MeOH, 80% MeOH, 1 hr each)
Wash in 100% MeOH 1 hr, 3 hr, O/N
Wash in 100% DCM 1 hr, 3 hr, O/N
Wash in 100% MeOH 1 hr, 3 hr, O/N
Rehydrate with reversed MeOH/B1n gradient (from 80% MeOH to 20% MeOH, 1 hr each)
Wash in B1n buffer: shake at RT, 1 hr, 3 hr, O/N
Wash with SdC buffer 4 hr, 24 hr x 2, 48 hr x 2
Wash in 5% DMSO/0.3M Glycine/PTwH 37°C, 6h, O/N
(Optional pause point, wash with PTwH 3 hr, 4 hr, O/N, then store in PTxwH, 4°C)
Incubate in primary antibody in 5% DMSO/0.3M Glycine/PTxH at 37°C O/N for 8 days
Wash with PTwH 3 hr, 4 hr, O/N
Wash with PTwH 24 hr x 4
Incubate in secondary antibody in PTxH at 37°C O/N for 8 days (protect from light from this point forward, secondary must have emission 560 or higher due to autofluorescence in 488 and below)
Wash with PTwH 3 hr, 4 hr, O/N
Wash with PTwH 24 hr x 4
Fix the sample with 2% PFA at 4°C O/N to stabilize the morphology.
Dehydrate in MeOH/H2O gradient (20% MeOH, 40% MeOH, 60% MeOH, 80% MeOH, 1 hr for each step)
Wash in 100% MeOH: shake at RT, 1 hr, O/N
Wash in 100% MeOH: shake at RT, 3 hr
Wash in 1:2 MeOH/DCM 2 hr, 100% DCM 1 hr, 3 hr, O/N
Wash in 100% DCM 1 hr, 1 hr, 3 hr, O/N
Clear in DBE 1hr, 4hr, O/N (DBE can melt regular plastic. EP tubes are fine. Use glass vials for long-term storage.)
Cleared sample can be stored in DBE in dark at RT or 4°C before imaging. (Fill the vial with DBE to minimize air oxidation for long term stability.)
LaVision BioTec Ultramicroscope is recommended for imaging
Depending on the age of rat, a custom holder may be needed to accommodate larger brain size
Mount the DBE-cleared sample onto the holder.
Load the holder to the imaging chamber filled with DBE
Set up imaging as instructed by LaVision user protocol
Sagittal or horizontal orientation is recommended as tissue will likely exceed lens working direction in coronal orientation
This whole mount immunolabeling protocol works well for sparse makers, but dense markers would be challenging to get complete staining. Most antibodies will be depleted on the superficial layers, leaving a ring-like pattern.
Double labeling can be done with two primary antibodies incubated simultaneously followed by simultaneous secondary antibodies
The exact time for each step in delipidation and others can be adjusted based on the property of the sample. Adipose tissue may need stronger extraction with one additional DCM step.
After DBE clearing and imaging, samples can be rehydrated back to PBS (through DCM, MeOH, the reversed MeOH gradient, and then PBS) for additional rounds of immunolabeling and analysis.
To reduce oxidation/discoloration, add 2-Mercaptoethanol (βME) at 120 uM (4.2 uL/500 mL) to MeOH steps
PBS (500 mL)
Perfusion
H20	 450 mL
PBS 10X	 50 mL
20% NaN3	 500 μL
Trim
PTwH (500 mL)
H20	 450 mL
PBS 10X	 50 mL
Triton X-100	 500 μL
Tween-20	 250μL
Heparin (20 mg/mL stock) 50 μL
20% NaN3	 500 μL
Delipidation
B1n (500 mL)
H20	 500 mL
Triton X-100	 500 μL
Glycine	 10 g
10M NaOH	 50 μL
20% NaN3	 500 μL
Mix until fully dissolved, store at RT
Immunolabeling
SdC (500 mL)
Clearing
H2O	 450 mL
PBS 10X	 50 mL
20% NaN3	 500 μL
methyl-β-cyclodextrin 5 g
Mix at RT until fully dissolved, filter and store at RT
Imaging
Additional Notes

### Slide 2
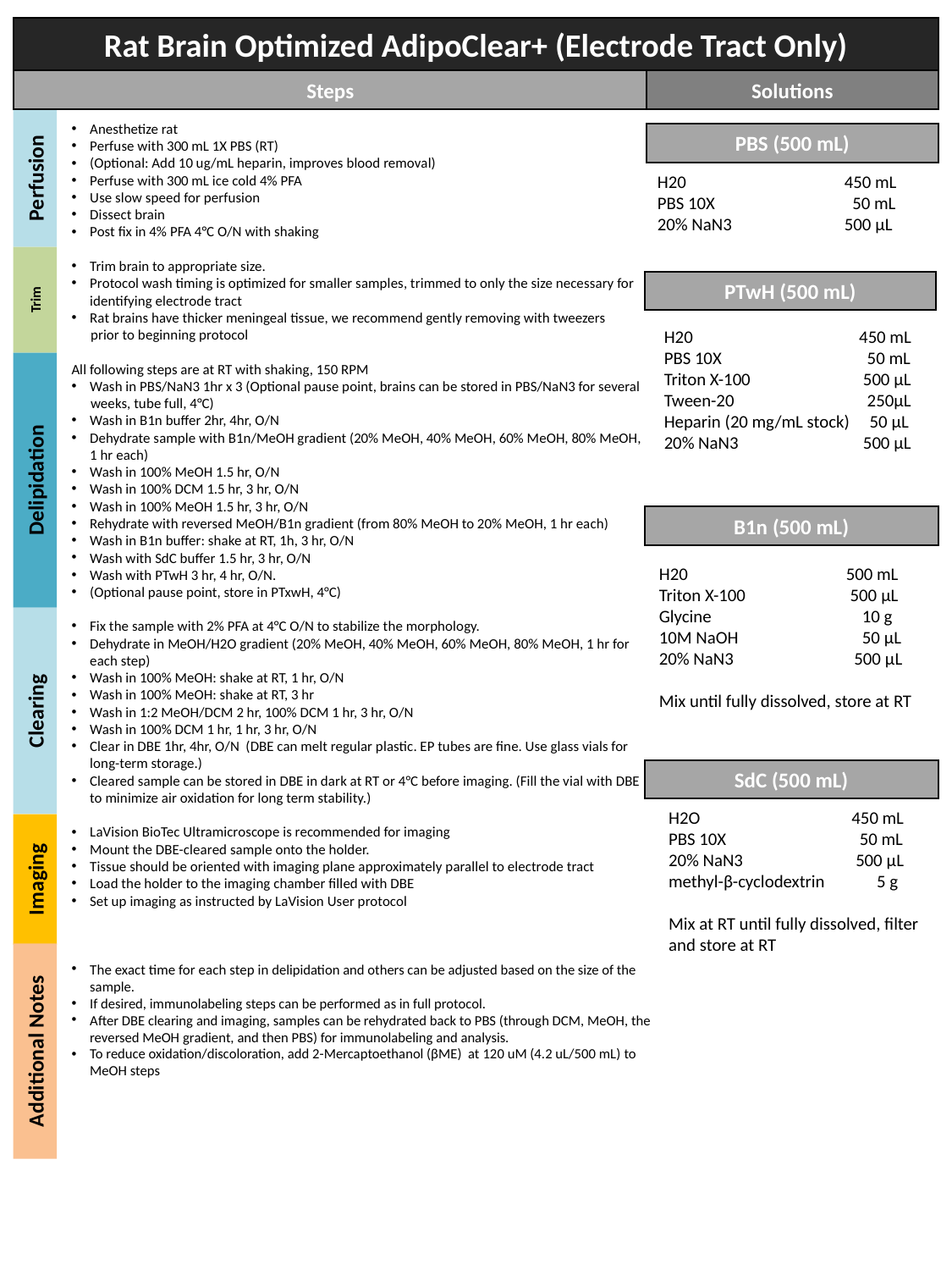

Rat Brain Optimized AdipoClear+ (Electrode Tract Only)
Steps
Solutions
Anesthetize rat
Perfuse with 300 mL 1X PBS (RT)
(Optional: Add 10 ug/mL heparin, improves blood removal)
Perfuse with 300 mL ice cold 4% PFA
Use slow speed for perfusion
Dissect brain
Post fix in 4% PFA 4°C O/N with shaking
Trim brain to appropriate size.
Protocol wash timing is optimized for smaller samples, trimmed to only the size necessary for identifying electrode tract
Rat brains have thicker meningeal tissue, we recommend gently removing with tweezers
 prior to beginning protocol
All following steps are at RT with shaking, 150 RPM
Wash in PBS/NaN3 1hr x 3 (Optional pause point, brains can be stored in PBS/NaN3 for several
 weeks, tube full, 4°C)
Wash in B1n buffer 2hr, 4hr, O/N
Dehydrate sample with B1n/MeOH gradient (20% MeOH, 40% MeOH, 60% MeOH, 80% MeOH, 1 hr each)
Wash in 100% MeOH 1.5 hr, O/N
Wash in 100% DCM 1.5 hr, 3 hr, O/N
Wash in 100% MeOH 1.5 hr, 3 hr, O/N
Rehydrate with reversed MeOH/B1n gradient (from 80% MeOH to 20% MeOH, 1 hr each)
Wash in B1n buffer: shake at RT, 1h, 3 hr, O/N
Wash with SdC buffer 1.5 hr, 3 hr, O/N
Wash with PTwH 3 hr, 4 hr, O/N.
(Optional pause point, store in PTxwH, 4°C)
Fix the sample with 2% PFA at 4°C O/N to stabilize the morphology.
Dehydrate in MeOH/H2O gradient (20% MeOH, 40% MeOH, 60% MeOH, 80% MeOH, 1 hr for each step)
Wash in 100% MeOH: shake at RT, 1 hr, O/N
Wash in 100% MeOH: shake at RT, 3 hr
Wash in 1:2 MeOH/DCM 2 hr, 100% DCM 1 hr, 3 hr, O/N
Wash in 100% DCM 1 hr, 1 hr, 3 hr, O/N
Clear in DBE 1hr, 4hr, O/N (DBE can melt regular plastic. EP tubes are fine. Use glass vials for long-term storage.)
Cleared sample can be stored in DBE in dark at RT or 4°C before imaging. (Fill the vial with DBE to minimize air oxidation for long term stability.)
LaVision BioTec Ultramicroscope is recommended for imaging
Mount the DBE-cleared sample onto the holder.
Tissue should be oriented with imaging plane approximately parallel to electrode tract
Load the holder to the imaging chamber filled with DBE
Set up imaging as instructed by LaVision User protocol
The exact time for each step in delipidation and others can be adjusted based on the size of the sample.
If desired, immunolabeling steps can be performed as in full protocol.
After DBE clearing and imaging, samples can be rehydrated back to PBS (through DCM, MeOH, the reversed MeOH gradient, and then PBS) for immunolabeling and analysis.
To reduce oxidation/discoloration, add 2-Mercaptoethanol (βME) at 120 uM (4.2 uL/500 mL) to MeOH steps
PBS (500 mL)
Perfusion
H20	 450 mL
PBS 10X	 50 mL
20% NaN3	 500 μL
PTwH (500 mL)
Trim
H20	 450 mL
PBS 10X	 50 mL
Triton X-100	 500 μL
Tween-20	 250μL
Heparin (20 mg/mL stock) 50 μL
20% NaN3	 500 μL
Delipidation
B1n (500 mL)
H20	 500 mL
Triton X-100	 500 μL
Glycine	 10 g
10M NaOH	 50 μL
20% NaN3	 500 μL
Mix until fully dissolved, store at RT
Clearing
SdC (500 mL)
H2O	 450 mL
PBS 10X	 50 mL
20% NaN3	 500 μL
methyl-β-cyclodextrin 5 g
Mix at RT until fully dissolved, filter and store at RT
Imaging
Additional Notes
